# Strawberry: fast and accurate genome-guided transcript reconstruction and quantification from RNA-seq

**DOI:** 10.1101/043802

**Authors:** Ruolin Liu, Julie A. Dickerson

## Abstract

We propose a novel method and computational tool, Strawberry, for transcript reconstruction and quantification from paired-end RNA-seq data under the guidance of genome alignment and independent of gene annotation. Strawberry achieves this through disentangling assembly and quantification in a sequential manner. The application of a fast flow network algorithm for assembly speeds up the construction of a parsimonious set of transcripts. The resulting reduced data representation improves the efficiency of expression-level quantification. Strawberry leverages the speed and accuracy of transcript assembly and quantification in such a way that processing 10 million simulated reads (after alignment) requires only 90 seconds using a single thread while achieving over 92% correlation with the ground truth, making it the state-of-the-art method. Strawberry outperforms Cufflinks and StringTie, the two other leading methods, in many aspects, including the number of corrected assembled transcripts and the correlation with the ground truth of simulated RNA-seq data.

**Availability:** Strawberry is written in C++11, and is available as open source software at https://github.com/ruolin/Strawberry under the GPLv3 license.

## 1 Background

Isoform-level quantification is a key step for differential alternative splicing and differential gene expression[25]. A number of software and computational methods have been developed for this [26, 19, 18, 3, 20]. However, most of the methods rely on existing gene annotations. This limits the use of such methods because even for the model organisms like drosophila melanogaster new isoforms are discovered all the time under different tissues and/or conditions [17]. [15] has shown that incomplete annotation is a major factor that negatively affects quantification accuracy for the existing methods. Thus, transcript-level quantification should be coupling with transcript assembly when dealing with RNA-seq data. Pure de novo assembly of raw RNA-seq is very challenging. Genome-guided methods like Cufflinks[26], instead, assemble aligned reads into transcripts. This allows them to take advantage of (if possible) a finished and high quality genome assembly and the-state-of-art spliced alignment algorithm.

Two strategies have evolved for tackling transcript assembly and quantification: simultaneously constructing transcripts and quantifying their expression or first constructing a set of transcripts then calculating their expression. Clearly, transcript reconstruction and quantification are closely related and many methods try to solve both simultaneously [3, 24, 16, 14] and this can be done in the framework of optimization or regularization. These methods usually exhaustively enumerate all possible transcripts and then use regularization to get rid of unlikely transcripts when calculating their expression. This occurs by punishing on the size of non-zero expression transcripts in favor of a smaller set of transcripts. Another strategy involves breaking the problem up in a step-by-step manner, like Cufflinks [26]. First, reconstruct a set of transcripts, then assume we know the set of transcripts and perform the quantification. Although it sounds like the all-in-one approach is more appealing, it is less computationally efficient than the step-by-step approach. Methods using all-in-one approach, e.g., Scripture [7] and iReckon [16] seek to enumerate all possible paths in order to reconstruct all isoforms expressed at a significant level. The authors of Cufflinks and Scripture call it a “maximum precision vs maximum sensitivity”problem [5]. However, the number of possible paths grows exponentially using the exhaustive principle. This makes it likely to generate more false positives impairing both accuracy and speed of abundance estimations. Therefore, to address the assembly problem methods that use maximum sensitivity have to simultaneously quantify the transcripts abundances to get rides of some unreal transcripts, i.e., those have no expression. Also, the all-in-one approach involves tuning the relative weight between the main objective and regularization term, which can vary between experiments.

### 1.1 Method overview

Strawberry adopts a step-by-step approach based on the previous arguments. The first step seeks a parsimonious representation of transcripts which best explains the input reads with the aid of flow network algorithms. Strawberry is different from the other methods which also utilize flow networks, FlipFlop [3], StringTie[19] and Traph[24], in that we only use flow networks to parsimoniously build up transcripts. The other methods use flow network to estimate the expression of transcripts, namely quantification. Traph uses min-cost flow to calculate isoform expression. StringTie uses a greedy algorithm to harvest the heaviest path and then uses maximum flow to estimate its expression. FlipFlop uses flow network to solve a regularized estimation problem as an all-in-one approach. Strawberry uses a flow algorithm to solve a parsimonious assembly problem, i.e., what are the minimum set of transcripts can best explain all the observed read alignments we observed. Compared to other three, Strawberry uses a “constrained” min-flow network which is specifically designed for paired-end reads. This requires a few modifications of canonical min-flow algorithm and to our knowledge Strawberry is the first to use the Constrained Minimum Path Cover algorithm(CMPC) to solve assembly problem.

After detecting the transcriptome, Strawberry calculates the pre-detected transcripts expression in RPKM and TPM using a generalized linear model (GLM) with Poisson density and identity link function which is based on the model of [22], Strawberry extends this model by grouping reads based on exon bins (see Methods section). This data reduction strategy greatly improves the computational efficiency with no or negligible sacrifice of performance. The intrinsic bias introduced from various RNA-seq steps also calls for a bias correction in the quantification. A recent paper from the authors of [22] proposed a way to estimate bias using a regularized approach[10]. We use, instead, a regression approach to which uses fewer parameters but is flexible enough to account for all sources of bias.

Strawberry can be used as a downstream analysis tool immediately after alignment, allowing it to take advantage of the latest genome assembly and state-of-the-art splice-awareness aligners. The mapping of paired-end reads may produce a fair amount of half-mapped reads. For those reads, Strawberry simulates the other end based on the mapped orientation and insert length distribution (if such information is not available Strawberry uses an empirical distribution inferred from the whole data set). In a nutshell, Strawberry accepts aligned RNA-seq data to construct transcript and estimates their abundances in a sequential manner(Fig.1). All of these can be done without gene annotations.

**Figure 1:**
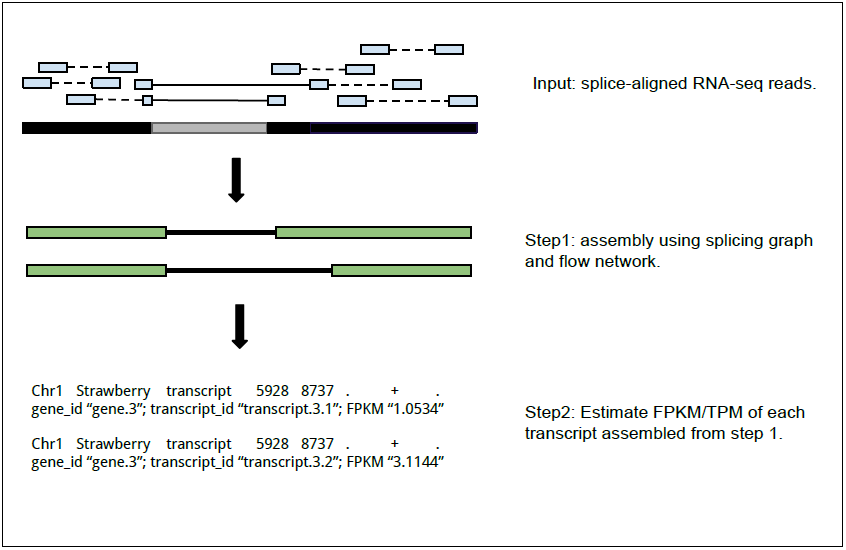
Method overview. It illustrates the input of this algorithm framework and its two major functions, in a step by step manner.

## 2 Results

### 2.1 Ground truth simulated data and programs to compare

We compare Strawberry to another two state-of-the-art program Cufflinks v2.2.1[26] and StringTie v1.2.0[19], on three simulation data sets, *RD25,RD60* and *RD100*. These data were generated by the procedure used in [15]—100bp paired-end reads generated only from 5800 multi-isoform Arabidopsis genes on TAIR10 genome version[12]. The only difference among these three data sets is the average sequencing depth. Roughly speaking, RD25 contains ~ 2.5*m*, RD60 ~ 6*m* and RD100 ~ 10*m* reads. And this procedure were repeated 10 times so that each data set consists of 10 RNA-seq libraries. Since plants genome have shorter intron than mammals, all the programs ran on the default parameters except for the maximum intron length, for which we set it to 5000*bp*. We chose Arabidopsis to simulate because it is a model organism and the complexity of its genome is between mammalian like Homo sapiens and simple organisms like C. elegans, making it a ideal test for separating the performance of the methods. Those simulated reads were then mapping against Arabidopsis TAIR10 genome assembly using Tophat2[11].

### 2.2 Comparing assembly accuracy

We use a Cufflinks module called Cuffcompare to compare the assembled transcripts or transfrags to the reference transcripts since those reads are all simulated based on the reference transcripts. For more details, see 7.2. We first compared the average number of matching transcripts, defined as the chain of introns between assembled transcript and the reference transcript complete match. Determination of transcription start and end sites is a known weakness of RNA-seq and impairs its application on identification of transcript boundaries [23]. Thus Cuffcompare defines a correct transcript as the chain of introns match with the reference, leaving possible variances in the first and last exon. We adopt this definition even for the simulated data. Fig. 5 shows that Strawberry consistently outperforms the others by giving more correct transcripts in all sequencing depth. We also observed that the differences among these three methods are well preserved under the three simulation data sets. This indicates the sequencing depth has a consistent impact on all the methods we are comparing and a higher read depth would usually lead to a better assembly result for the genome-guide method according to this simulation. Across all the 30 simulated RNA-seq libraries, Strawberry produced 1143 more accurate transcripts than StringTie did and 546 more than Cufflinks did.

**Figure 2:**
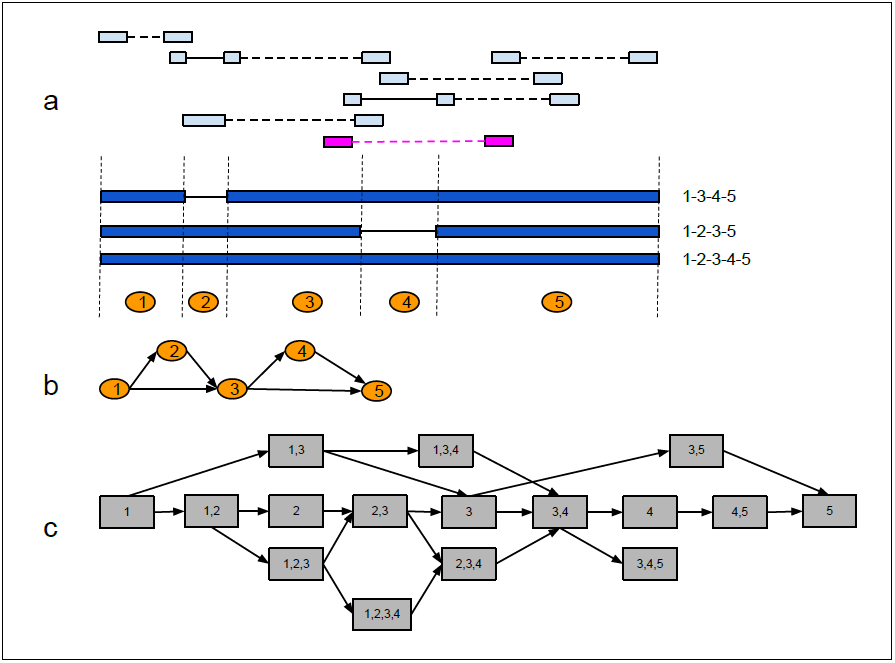
2(a), translation of read alignments into a splice graph. Seven imaginary paired-end reads (or fragments) are represented by light blue boxes intersected by solid lines (splicing junctions or introns) and broken lines (fragment inner distance) on the top; the arrow represents transcription direction. 3 likely isoforms reconstructed from the 7 alignments are shown below. Exons are shown as dark blue boxes and introns are shown as solid lines. 5 exon pieces (numbered solid circle) are created through breaking the biological exons. 2(b), the splicing graph constructed from part (a). Arrows indicate the transcription direction. 2(c), the exon bin map from part (b). An exon bin consists of one or more exon pieces covered by an entire read fragment. There is an directed edge connecting two bins, *x* → *y*, only if exactly one of the following conditions holds for true: i), *x* is the prefix of *y*. ii), *y* is the suffix of *x*. And iii), there is a direct edge from *x* to *y* in the original splicing graph and no single read span them. For high throughput experiment like RNA-seq, the last situation could be fairly uncommon. The magenta fragment can be distributed to two exon bins: 3-5 and 3-4-5. It is in consistent with all of the three isoforms.

**Figure 3:**
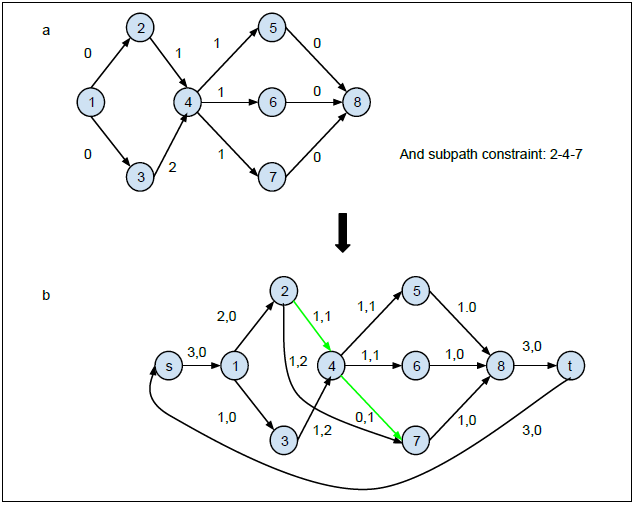
3(a), an input flow network with a subpath constraint {2-4-7}. The number next to an edge is the edge cost. For every edge *e*, the edge constraint imply 1 ≤ f(e) ≤ inf. 3(b), the transformed min-flow circulation network. The numbers (a,b) next each edge indicate the optimal flow on the edge and the edge cost respectively. After Step 3 the path constraints set *P*^*sub*^ = {(1,2), (1,3), (2, 4, 7), (4, 5), (4, 6), (5,8), (6, 8), (7, 8)}. Two edges are no longer in the constraint set are shown in green color. For these two edges, the minimum flow requirement is 0; for the rest of edges it is 1. Two dummy node *s* and *t* are added to complete the circulation. The number of flow after decomposition equals to the min flow which is 3.

**Figure 4:**
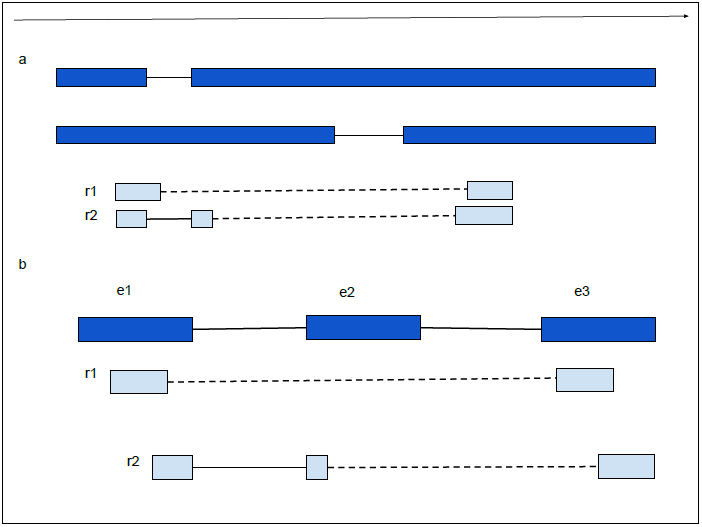
4(a), a pair of reads (light blue) have the same farthermost 5’ and 3’end but belong to different transcripts (dark blue). Dashed line represents the gap and solid line stands for spliced alignment. 4(b), read I belongs to exon bin {el,e3}; read 2 belongs to exon bin {el,e2,e3}

**Figure 5:**
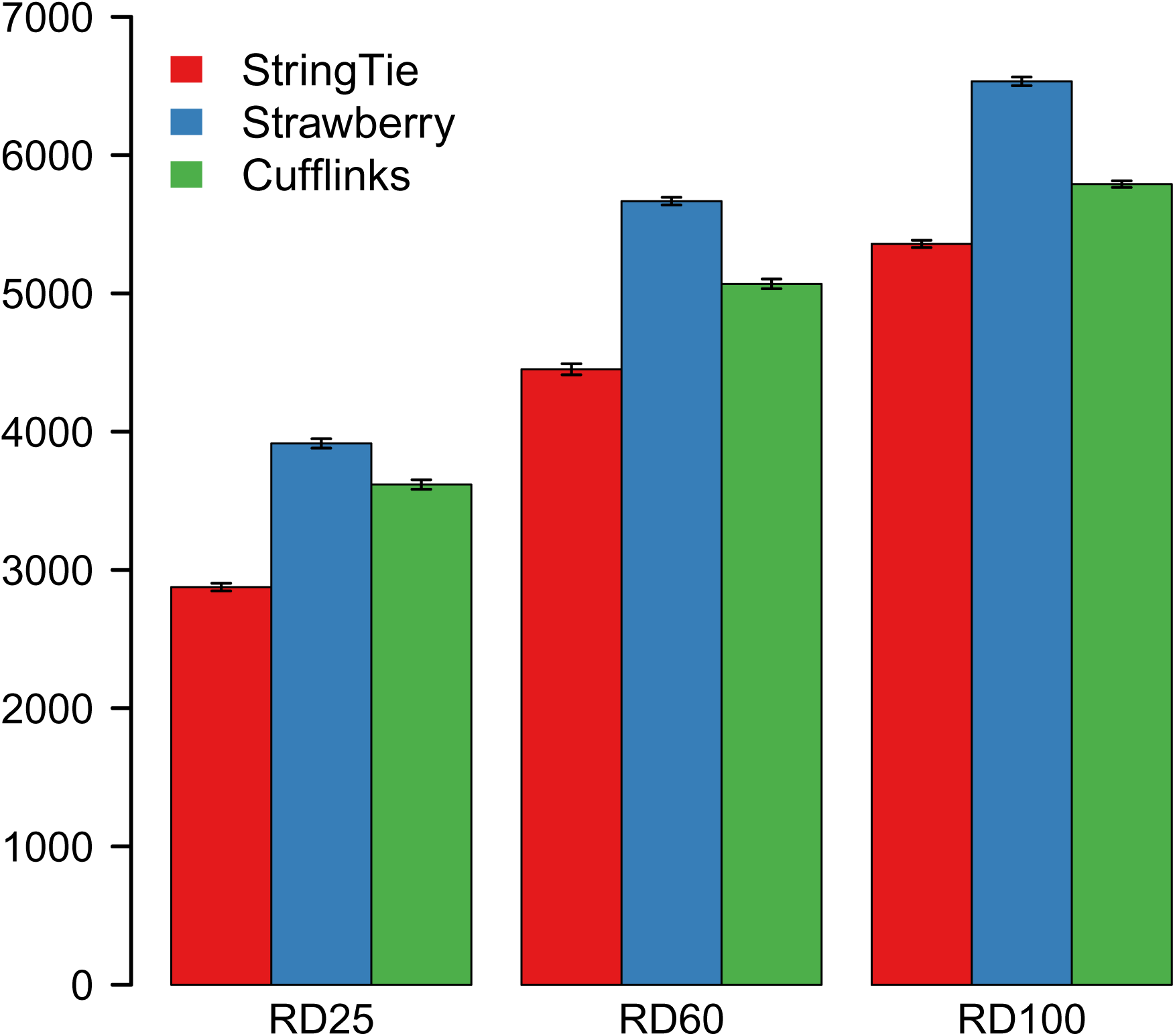
Number of correct transcripts given by each method under three different simulation data sets. Each data set consists of 10 replicates so the error bars are shown on top.

We then assessed the degree to which transcripts reported by each method matched the reference annotation at nucleotide, exon, intron and transcript level for three different sequencing depths (Fig. 6, 7 and 8). In all comparisons, Strawberry has a clear advantage against the other methods over sensitivity while maintaining roughly the same level of specificity. In RD100 data, for example, Strawberry clearly does better than the other two method by averaging 69.92%, 78.13%, 48.25% on sensitivity at exons, intron, and full transcripts level respectively, followed by Cufflinks, 65.51%, 74.09%, 42.76% and then StringTie, 63.44%, 73.47%, 39.58%. For all the methods, the sensitivity decreased as sequencing depth decreases while the specificity remains at a high level and doesn't change much. This indicates that although lower read depths make it harder for these methods to recover the true signal the results are still very reliable. Correct detection of full transcripts using RNA-seq data is still a very challenging task for all assemblers. Given sufficient sequencing depth (RD100), all methods can correctly retrieve 65%+ exons, 75%+ intron but no more than 50% full transcripts. On the other hand, specificity for exons and intron detection are also very high for all methods, especially for Strawberry, both averaging 99%. However, for transcript detection, this number comes down to 77% despite the fact that it is still the best among these three. For the methods that parsimoniously assemble reads into transcripts, this hints some rooms for improving—although the individual exons and introns are correctly recovered the ways to stitch them together are still not optimal.

**Figure 6:**
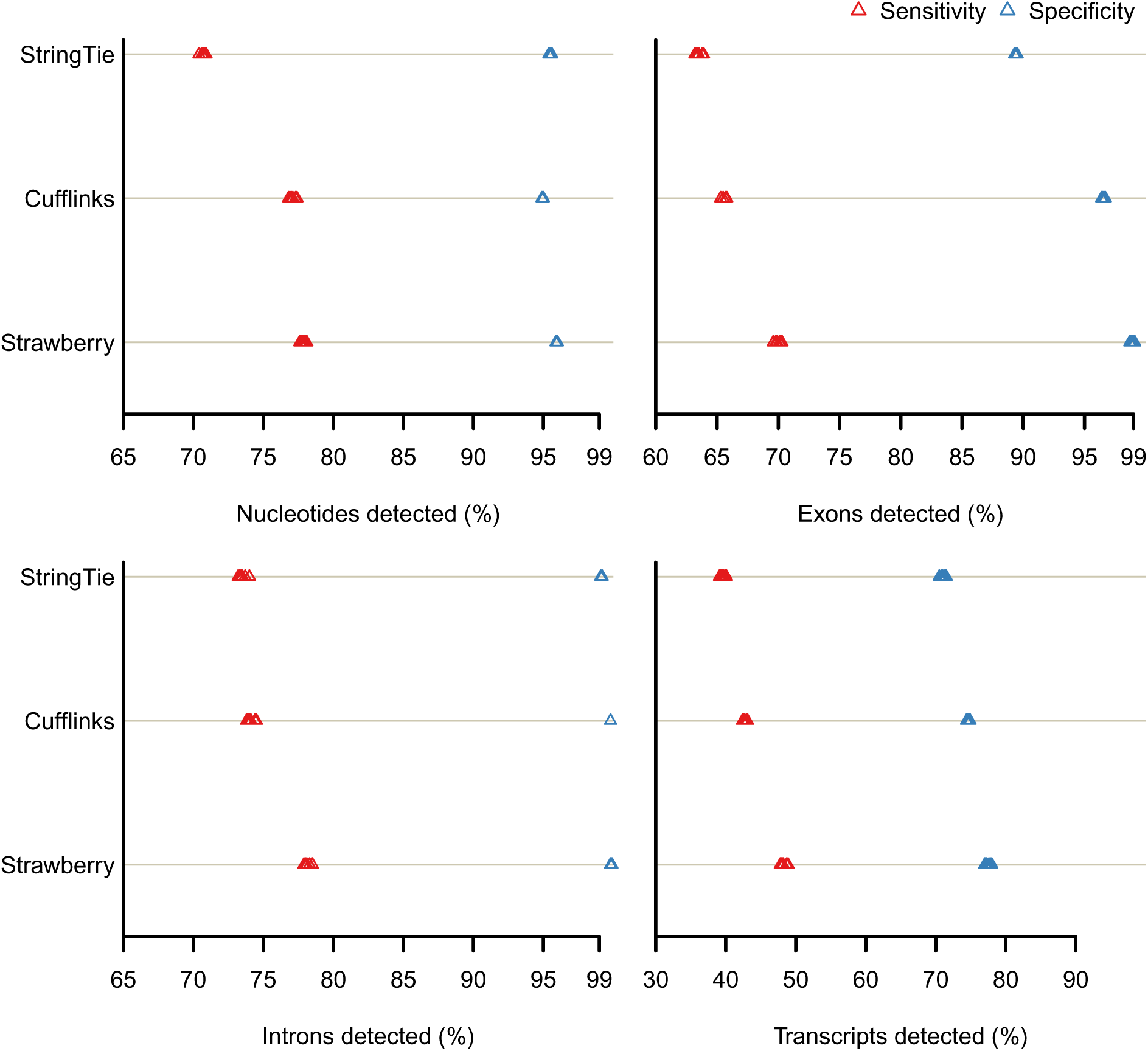
RD100

**Figure 7:**
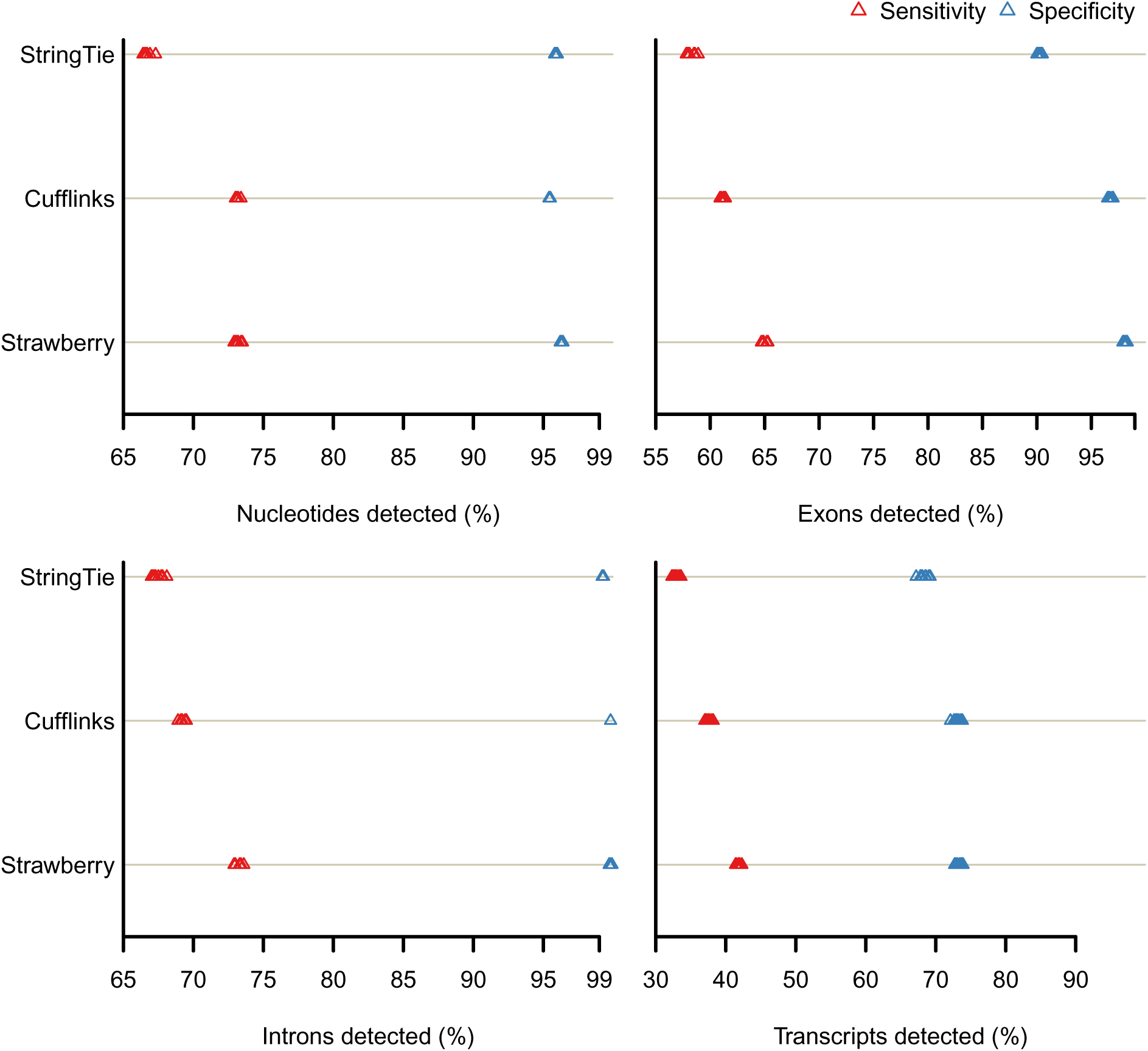
RD60

**Figure 8:**
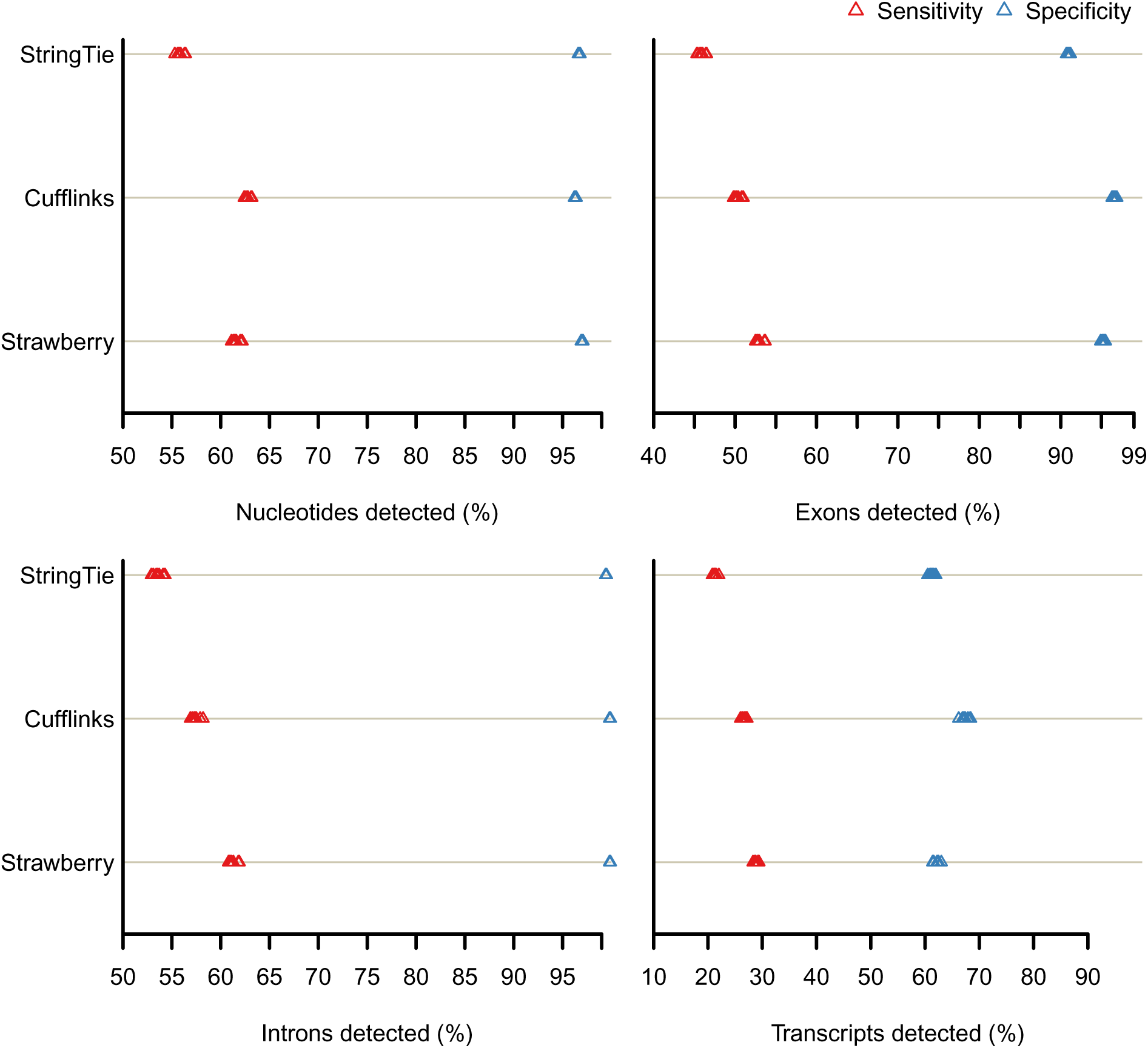
RD25

### 2.3 Comparing quantification accuracy

We adopt the metrics used in [18].

1. Proportionality correlation

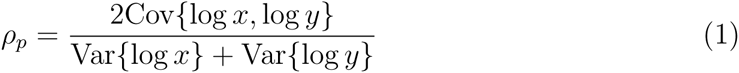

where *x*_*i*_ is the true value of the FPKM for isoform *i* based on ground truth simulated data and *y*_*i*_ is the predicted value. FPKM (Fragment per kilobase per million mapped read) is defined as 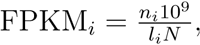, where *n*_*i*_ is the number of reads belonging to isoform *i*, *l*_*i*_ is the length of isoform *i*, and *N* is the total number of reads. FPKM values were incremented by 1 before log transformation to avoid infinite numbers.
2. Spearman correlation of between the true FPKM values and predicted FPKM values.
3. Mean Absolute Relative Difference (MARD), which is computed using the absolute relative difference ARD_*i*_ for each isoform *i*:

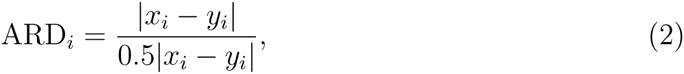

MARD is the mean value of the {ARD_*i*_|*i* ∈ 1,…, *I*}. ARD is bounded above by 2 and below by 0 and smaller value indicates a better prediction. [18] computes MARD on the number of reads deriving from each transcript which is commensurable to FPKM values.

Fig. 9, 10, 11 show the histogram of the three measures over 10 replicates for all three read depth data sets respectively. In these simulations, It can be seen that these methods are all well separated in terms of the all evaluation metrics except for only one case in which StringTie and Cufflinks are virtually tied over Spearman correlation in RD60 data (see Fig. 10). In the best performing case (RD100 data), Strawberry averaged 0.93, 0.927, and 0.377 on Proportional correlation, Spearman correlation and MARD respectively, followed by StringTie, 0.868, 0.859, 0.424 and then Cufflinks, 0.834, 0.876, 0.450. Cufflinks actually out-performs StringTie under Spearman correlation. Like the assembly results, the sequencing depth seems to have a uniform impact on the quantification accuracy and all methods favor the highest read depth. It worth mentioning that our enumeration of sequencing depth has not covered the full possible range. If we did we would expect a plateau or even a backfire, i.e., sequencing depth too high might actually hurts quantification. Overall, Strawberry out-performs the other methods under all evaluation metrics and sequencing depth and StringTie performs better than Cufflinks. However, the distance between the second and third place is less than distance between the first and second place. We also observe that Strawberry and StringTie have less variability in results than Cufflinks did, suggesting they are more consistent in terms of their estimates.

**Figure 9:**
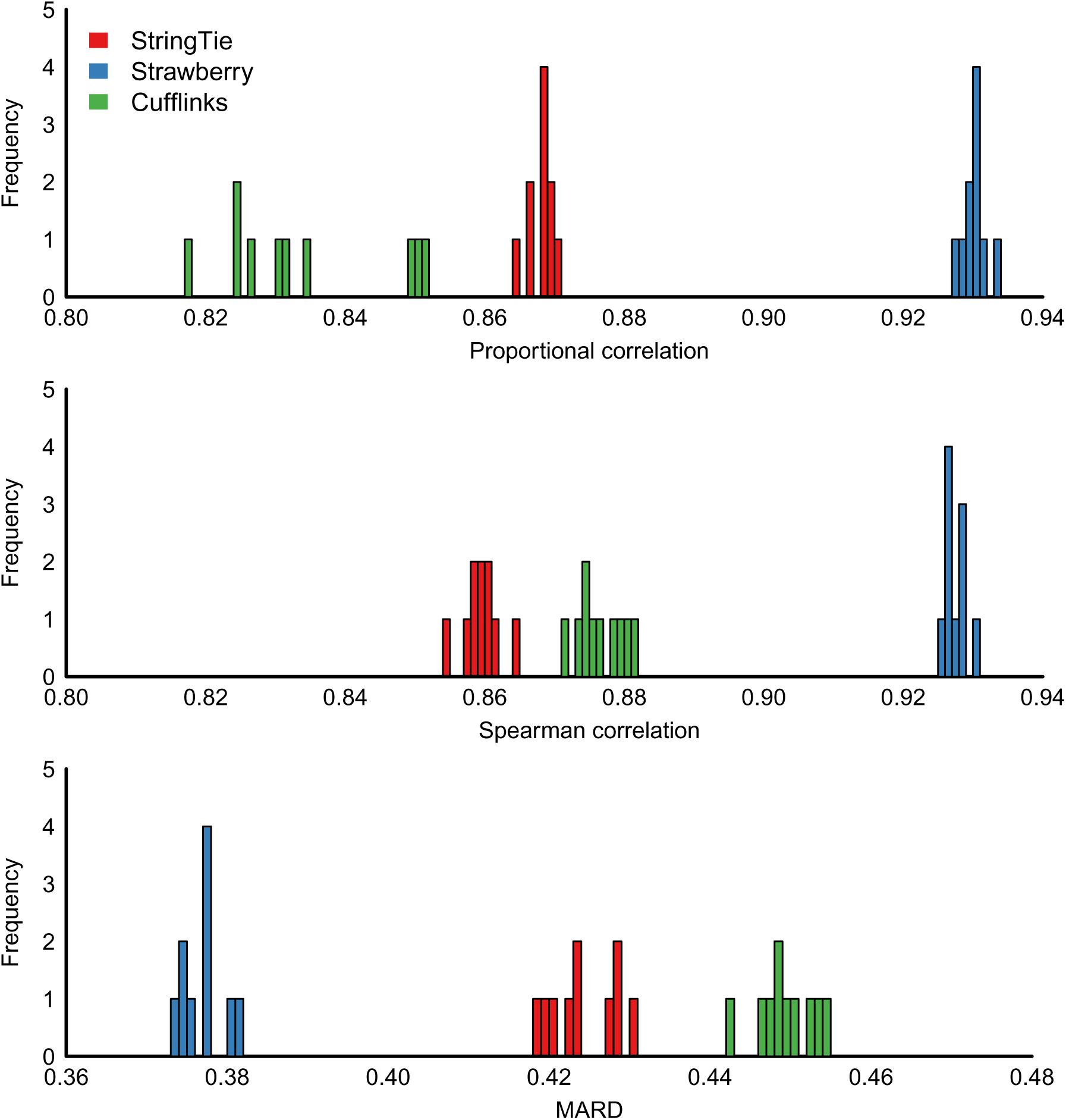
RD100

**Figure 10:**
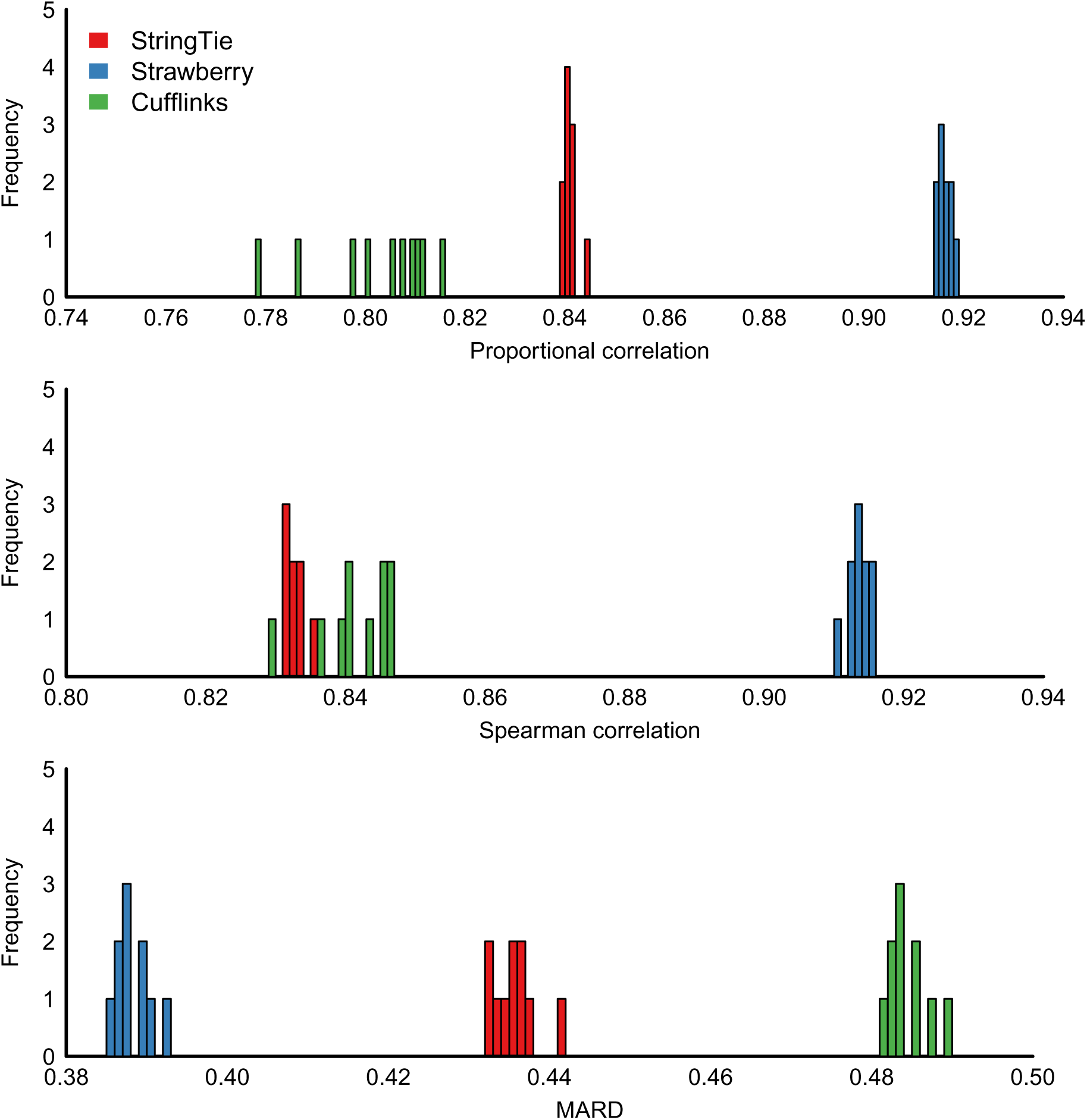
RD60

**Figure 11:**
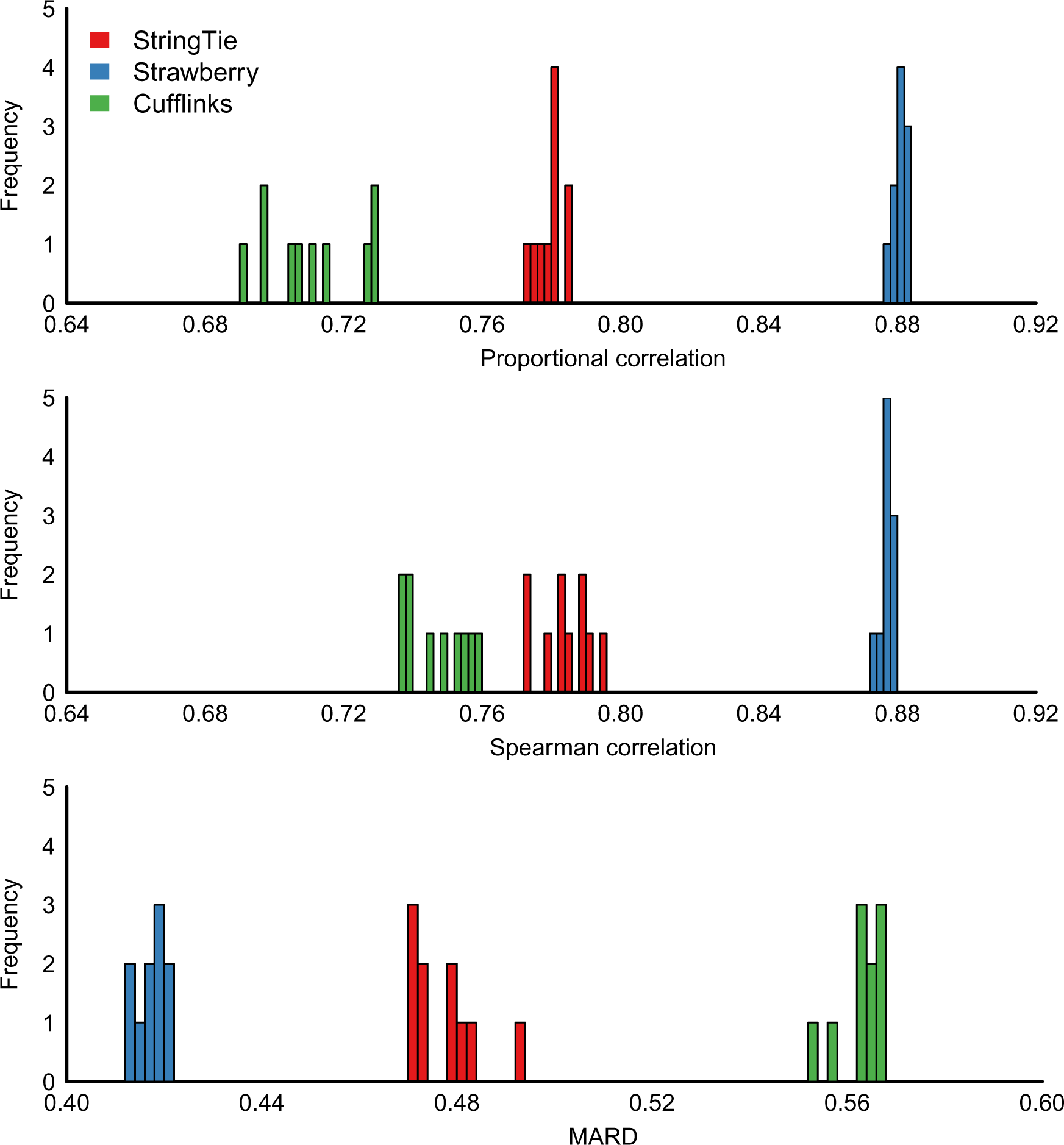
RD25

### 2.4 Real RNA-seq data

To demonstrate Strawberry utility on real data, we tested all three programs on the Homo sapiens HepG2 data from *Steijger et* al.[23]. The data was download from http://www.ebi.ac.uk/arrayexpress/experiments/E-MTAB-1730/, which includes alignment results from a library of 100 million 76bp paired-end Homo sapiens RNA-seq reads and a total of 140 NanoString probe counts. These 140 probes targeted 109 genes, designed against specific isoforms. NanoString counts were then compared to the highest RPKM value reported for isoforms consistent with the probe design [23]. We follow the same procedure used in *Steijger et al*. excepts for using Tophat2 alignment result and Cuffcompare for finding the best matching transcripts. Pearson correlations between the log-transformed RPKM and NanoString count and the number of probes having matched transcripts were very close for all three programs. Strawberry had the edge over the other two by only 1% (Table 1). It is probably because this is such a small dataset. More than one third of the probes have no matching transcripts and subsequently an PRPKM of 0 were assigned. If we applied the same Pearson correlation metric to probes with matching transcripts only, Strawberry became a more clear front-runner(Table 1).

**Table 1:** Quantification results on real RNA-seq data HepG2. NanoString counts from 140 probes were compared to the RPKM values reported for three programs. If no such transcript was reported by programs for a designed probe, an RPKM of 0 was assigned in the all-probes comparison (first line) or was simply excluded in the matched-probes comparison (second line). The number of probes which have matching transcripts is reported on the last line.

|  | Strawberry | Cufflinks | StringTie |
| --- | --- | --- | --- |
| Pearson corr. for all probes | 0.681 | 0.672 | 0.669 |
| Pearson corr. for matched probes | 0.500 | 0.458 | 0.455 |
| # probes with matching transcripts | 82 | 82 | 83 |

It's worth mentioning that the numbers reported here are not directly comparable to the numbers in *Steijger et al*. because of the two differences in the analysis pipeline aforementioned. However they are pretty close. The Pearson correlation for all probes of Cufflinks is 0.68, which is the highest in *Steijger et al.*.

### 2.5 Running Time

The above programs were tested on a Dell Precision T1650, equipped with Intel Core i7-3770 CPU and 16GB RAM. The running times of all programs using a single thread on a sample from the RD100 data set(approximately 10 million reads) were given in Table 2. Each program was given the aligned data in BAM format and the time spent on alignment is not included. The simplicity of StringTie algorithm makes it the fastest method among the three. Cufflinks and Strawberry both use EM algorithm for ambiguously assigning reads to transcripts. The EM algorithm is a time consuming algorithm but the reduced data representation by Strawberry makes it almost 10 times faster than Cufflinks.

**Table 2:** Total Runing time in minutes for processing one sample on RD100 data set

| StringTie | Strawberry | Cufflinks |
| --- | --- | --- |
| 0.5m | 1.48m | 10.32m |

## 3 Discussion

Strawberry adopts a step-by-step approach for transcripts assembly and quantification of expression levels. The assembly and quantification are an inseparable whole in a since that every eukaryotic RNA-seq experiment is likely to generate unknown transcripts even for the well-annotated species, let along the fact that gene models for many organisms are under-developed. And our previous study of alternative splicing has shown that a incomplete genome annotation can have a huge negative impact on the quantification accuracy[15]. Strawberry avoids using gene annotations but it is not a purely de novo assembly method. Strawberry's transcriptome assembly takes advantage of the latest genome assembly and state-of-art splice-awareness aligners and is usually more accurate then the de novo assemblers. However, this makes Strawberry relied on alignment results, adding its dependency on alignment algorithms. We often experience different results based on different alignment packages.

Both Cufflinks and Strawberry use EM algorithm for optimizing the likelihood functions. However, because of a reduced data representation, Strawberry is 10 times or more faster than Cufflinks. StringTie uses a flow algorithm for quantification which is very fast compared to the EM algorithm that Strawberry and Cufflinks use. This makes it unlikely for Strawberry to outrun StringTie. The result of computing time in the last section was based on a single thread. Like StringTie and Cufflinks, Strawberry implements the thread-level parallellism which can process several locus at a time to greatly speed up the program.

The lack of golden standard data for assessment of RNA-seq applications is still a major problem for the community. The comparison use in this paper is primarily based on simulated data where we know the ground truth. However, the simulation programs can fall short in various aspects to resemble real data including sequencing bias, read errors, etc. Interestingly, using bias correction in Strawberry and Cufflinks does not lead to an increase in performance using the simulated data. StringTie does not have the option to allow user to enable/disable its bias correction and therefore we can not judge. The same pattern was also observed when using HepG2 data. This is probably because NanoString is not necessarily a more reliable technology compared to RNA-seq.

## 4 Conclusion

In this paper, we have introduced Strawberry, a fast, accurate genome-guide assembler and quantification tool for RNA-seq data. It facilitates discovery of novel isoforms and calculation of transcript-level expression. Based on our simulation, Strawberry not only recovers more true transcripts while achieving the same false discovery rate in assembly compared to two other leading methods but also outperforms them on the quantification accuracy. The other advantage of Strawberry is its speed. It finished processing 10 million simulated reads (after alignment) within 90 seconds on a 16GB RAM and core-i7 desktop using a single thread, 10 times faster than Cufflinks. Strawberry achieves this level of speed and accuracy through applying min-cost min-flow algorithm to assembly, a reduced data representation to exon bin counts and a model-based approach to adjust sequencing bias and to estimate transcript abundances simultaneously.

## 5 Methods

### 5.1 Assembly problem formulation

Two computationally alternative strategies have emerged for transcriptome reconstruction. Genome-independent methods assemble transcripts directly from RNA-seq reads independent of any reference genome. This is important when the reference genome is not available, gapped, highly fragmented or largely altered, e.g., in cancer cells [6]. Genome-guided methods assemble, instead, aligned RNA-seq reads (usually in BAM/SAM format) into transcripts. The alignment relies on a reference genome (if possible a finished and high quality genome) which usually leads to a better assembly. Cufflinks is one of the pioneers that use mapped reads to assemble transcripts [26]. We here summarize the Cufflinks assembly algorithm using an optimization language. Given the aligned reads on one locus, the set of *n* fragments *R* = {*r*_*i*_ : *i* ∈ 1,…,*n*} form a partially ordered set in which *r*_*i*_ ≤ *r*_*j*_ if and only if the start position, in the transcription direction, of *r*_*i*_ is less than or equal to *r*_*j*_ and the two are compatible. In brief, two fragments are incompatible if they imply different introns. A more comprehensive definition of incompatibility is given in [26]. We define a path p as an ordered set of fragments {*r*_0_,…, *r*_*i*_,…, *r*_*n*_}. Thus the assembly problem can be expressed as finding a path cover *C*, defined as

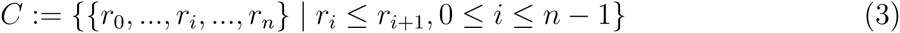

which minimizes

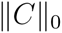

and satisfies

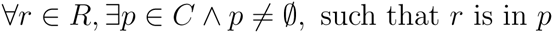

This means that we want to find a minimum number of paths that cover every read at least once. And due to the help of alignment, reads are well separated into loci and each locus can be processed independently.

The problem can be simplified using splicing graphs [9], that are intensively used in genome-guided methods. Splicing graph *G* = (*V*, *E*) is defined as a directed acyclic graph (DAG) on the set of transcribed positions *V* and edge set *E*. *G* contains a edge from *v*_*i*_ to *v*_*j*_ if and only if *v*_*i*_ and *v*_*j*_ (*v*_*i*_ ≠ *v*_*j*_, for *i* ≠ *j*) are consecutive positions in at least one transcript. It is common to refine the graph *G* by collapsing consecutive vertices if all of them have only one outgoing edge and one ingoing edge. When doing so, the vertices *V* of graph become exons (or partial exons) and edges E become introns [9]. For convenience, we use exon uniformly. Another common modification is adding an artificial source vertex and target vertex, where the source vertex, denoted by *v*_*s*_, connects to each of the smallest position of transcripts and the target vertex, *v*_*t*_, connects to each of the largest position of transcripts.

Strawberry uses splicing graph representation and instead of assembling reads it assembles exons into transcripts. This make the assembly algorithm more time efficient without sacrificing much of the performance since all reads on a exon belong to the same set of transcripts. This exon representation also has a positive impact on the quantification part. The read counts on exons can be seen as sufficient statistics for our quantification model. Strawberry also associates with each exon a bias factor to account for various sources of bias that affect the distribution of read coverage along transcripts, which is generally considered as nonuniform [21]. The idea of quantification and bias correction is discussed in more detail in the quantification section.

Under the splicing graph representation, the assembly problem is reduced to finding a minimum set of exon paths. We add additional constraints that each exon path has to start from source node *v*_*s*_ and end at target node *v*_*t*_. These two nodes are not real exons and are added to make use of the flow algorithm. However, these nodes introduce a sound intuition for transcript assembly using the flow algorithm: each transcript is an ordered sequence of exons from a promoter region to terminator. Therefore an exon path is defined as a directed path from *v*_*s*_ to *v*_*t*_, so called (*s*, *t*)-path. Also, it highlights a limitation in the proposed method as it assumes that all transcripts have the same promoter region, which might not be true in some situations.

### 5.2 Constructing a weighted splicing graph

A Splicing Graph, equivalent to a DAG, is used to represent the set of transcripts at any identified locus. The non-overlapping loci are treated independently to allow for maximum parallelization. The continuous stretch of read coverage without gaps are retrieved first as primitive exons and junction alignments are retrieved to form introns. These introns are filtered according to how many reads support them and the attributes of these reads. The remaining introns are then used to refine these primitive exons to form the splicing graph(Fig. 2a). We use the same criteria to discard an intron as in [26]. The thresholds are arbitrary but work well in practice.

- more than 70% of the reads are not uniquely aligned.
- If two introns overlap and one’s expression is less than 5% of the other, then the lower one is removed. The intron expression is calculated by the total junction reads.
- The number of small overhang reads supporting a junction is likely to be small under the assumption that reads are distributed uniformly along the their parent transcripts. The small overhang read is a particular junction read defined as one end of the read is mapped within a small distance (e.g. 6 bp) of exon-intron boundaries. The expected number of small overhang reads is calculated by a binomial distribution, *Bin*(*n*,*p*), where *n* is total junction reads and 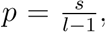, *s* being the small overhang distance and *l* being read length. When *n* is large (e.g., > 100), we use a normal approximation as *N*(*np*, *np*(1 – *p*)).

Next, Strawberry defines the initial exon blocks as a continuous region covered by read. Then the initial exon blocks are divided into smaller pieces if they are interrupted by filtered introns. Therefore, each exon piece (for convenience we still call it exon) is either fully contained or excluded in one or more transcripts. In the DAG representation, there is an edge between two exons if two exons are connected by intron or two exons are consecutive in their genomic coordinates (see Fig.2(a)). The number of reads spanning two exons is used as edge weight as it shows the belief of two exons are in the same transcript. For the intron edges, the weight is simply the total junction read number. In the implementation,a negative sign is added since the algorithm solves for the minimum weight.

### 5.3 Optimization with flow network

The splicing graph formulation allows us to consider the assembly problem as finding a set of (*s*, *t*)-paths on graph *G*. This is related to a canonical computer science problem known as the Minimum Path Cover (MPC) problem.

#### Definition 1

MPC problem. Given a directed graph G = (V, E), a path cover is a set of directed paths such that every vertex v ∈ V belongs to at least one path.

The ordinary MPC problem is not a perfect fit for the splicing graph since it only requires that every exon shows up at least once. This leaves the possibility that some introns might not be covered. Also, if paired-end reads are used, it is likely that they span more than two consecutive nodes. These nodes constitute a subpath that also must be covered by at least one final path. These constitute the constraints from the mapped reads. A efficient algorithm for the Constrained MPC (CMPC) problem has been advanced in [20]. Cufflinks uses an overlap graph where read alignments are considered as nodes and edges stand for compatibility between reads. The incompatible reads are left out [26]. It is, in fact, a MPC problem applied to the individual reads.

Following the terminology in [20], the CMPC problem is formulated in the following definition.

#### Definition 2

CMPC problem. Given a DAG G with nodes V(G) and edges E(G), and a weight w(e) for each e ∈ E (G), and a subpath family 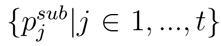 the task is to find a minimum number of k of directed paths {p_i_|i ∈ 1,…,k} in G such that

- Every node in V(G) occurs at least once in some p_i_.
- Every edge in E(G) occurs at least once in some p_i_.
- Every path 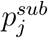 is entirely contained in some p_i_.
- Every path p_i_ starts in S and ends in T.
- 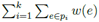 minimum among all solutions of k paths.

It has been shown in [20] that CMPC problem can be reduced to the canonical MPC problem with only vertex constraints. The canonical MPC problem can be solved using one of the well established flow network algorithms, e.g., the min-cost circulation flow algorithm [1], Therefore a strong polynomial time solution is guaranteed.

#### Definition 3

Flow network is a DAG G = (V,E) with source v_s_ ∈ V(G) and target v_t_ ∈ V(G), where edge e ∈ E(G) has capacity upper and lower limit l(e), u(e) and flow /(e) with the following three properties.

- Capacity constraints: l(e) ≤ f(e) ≤ u(e), for all e ∈ E(G).
- Flow conservation constraints: Σ_e=(uv)_f(e) = Σ_e=(v,u)_f(e), for all v ≠ v_s_, v_t_. This means the incoming flow is equal to outgoing flow.

Generally speaking, to solve a flow network problem is to construct a map, *f* : *E* → *R*, which maps an edge to a real number, or most commonly an integer number. This number is called a flow. The flow decomposition theorem (see, e.g. [1]) guarantees the flow network can be used to solve MPC problem. It says that for flow *f*(*e*) on edge e can be decomposed into a set of flows on (*s*, *t*)-path. However the decomposition is not unique. To overcome this, we adopt a greedy algorithm to decompose the result from flow algorithm. These all together consist the algorithm solving the CMPC problem.

#### Algorithm 1

Constrained Minimum Path Cover algorithm (CMPC)

1. Let 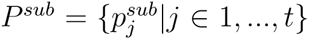 denote the set of subpath constraints. Then add all edges to P^sub^, that is P^sub^ ∪ e for every e ≨ E(G).
2. For every pair of path constraints 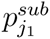 and 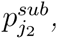, drop the duplicates, that is 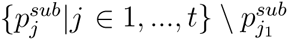, if 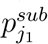 is contained in 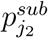.
3. For every path constraint 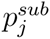 which starts at node u and ends at node v, do:

- E (G) := E (G) ≪ {(u,v)}
- *Set the lower and upper bounds for flow form u to v*. *lower*(*u*,*v*) = 1 *and upper* (*u*, *v*) = inf
- 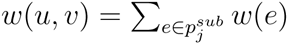
4. *For each e ∈ E(G) and e ∉ P^sub^, set lower(e) = 0 and upper(e)* = inf.
5. *Add an edge (t,s) from sink node t to start node s to complete the circle. Set lower and upper bounds for this edge as well. lower(t, s)* = 0 *and upper*(*t, s*) = inf.
6. Compute a min-weight min-flow circulation on this transformed input G with the following properties.

- G is a flow network which satisfies capacity constraints and flow conservation constraints.
- Min flow: Σ_e∈E(G)_ f(e) is minimum.
- Min weight: Σ_e∈E(G)_ w(e) is minimum.
7. Finally, the flow on edge (v_t_,v_s_) equals to the min flow and minimum number of (s,t)-paths. We decompose the flow network into this number of paths and each path corresponds to an assembled transcript.

Fig.3 demonstrates a toy example of this algorithm.

### 5.4 A GLM Poisson model for quantification

Strawberry’s quantification model is based on the insert length model proposed in [22] so we have kept the same notation whenever possible. Like the assembly, this model is also designed for a single locus from a single sample so that each locus is treated independently to allow for maximum parallelization. A summary of notation used is given int table 3. Strawberry extends this model by collapsing reads from same exon bins from the splicing graph. This change greatly reduces the number of observations and speed up the calculation. It also requires an adaptation for the definition of read type. A read type *s*_*j*_ in [22] refers to a specific position (which can be denoted as the 5’ end of the read) for single-end reads or two specific position (5’ end of the first read and the 3’ end of the second read) in paired end sequencing. The read type *s*_*j*_ in Strawberry is defined as two specific positions for a single end read (5’end and 3’end of the single read) and four specific positions for a paired end read (5’end and 3’end for the first read and then 5’ end and 3’end for the second read). This change is made because later reads are grouped by the exons they cross and having only two farthermost positions the same does not guarantee the same type reads have the same transcripts origin due to the gap alignment (Fig. 4(a)). Further reads are grouped into exon bins, based on those 4 positions (Fig.4(b)).

**Table 3:** Summary of notations

| Symbol | Meaning |
| --- | --- |
| $I$ | Total number of unique transcripts for a given locus in the sample, $i$ for indexes. |
| $J$ | Total number of unique reads for a given locus, $j$ for indexes. |
| $K$ | Total number of exon bins for a given locus, $k$ for indexes. |
| $\theta_i$ | The expected number of reads generated from $i$th transcript, $i = 1, \dots, I$. Those are the parameters to be estimated. |
| $\theta$ | The isoform abundance vector $[\theta_1, \dots, \theta_I]$ |
| $s_j$ | Read type $j$, $j = 1, \dots, J$. |
| $e_k$ | Exon bin $k$, $k = 1, \dots, K$. |
| $n_{i,j}$ | The number of reads $s_j$ that are generated from transcript $i$. |
| $n_j$ | The number of read $s_j$ that are generated from all the transcripts for a given locus. |
| $n_{i,k}$ | The number of reads overlapping exon bin $k$ that are generated from transcript $i$. |
| $n_k$ | The number of reads falling on exon bin $k$. |
| $n$ | The exon bin count vector. $[n_1, \dots, n_k]$. |
| $a_{i,k}$ | Up to a normalizing constant the sampling rate of $n_{i,k}$, that is, the rate that read falling on exon bin $k$ is generated from each individual transcript $i$. |
| $a_k$ | The sampling rate vector $[a_{1,k}, a_{2,k}, \dots, a_{I,k}]$ |
| $l(i)$ | The length of $i$th transcript. |
| $l(k)$ | The length of $k$th exon bin. |
| $l_{i,j}$ | The insert length of read type $s_j$ under isoform $i$. |
| $l_{ik}(j)$ | Number of possible positions that a read type $s_j$ can be generated on the $i$th isoform and $k$th exon bin. |
| $m$ | The total number of reads for a given locus in the sample. |
| $q(l_{i,j})$ | Empirical insert length distribution for read type $j$ and isoform $i$. |
| $t(\psi_{i,j})$ | Start site (5'end) distribution for read type $j$ and isoform $i$ at site $\psi_{i,j}$. |
| $b_k$ | The bias factor for exon bin $k$. |
| $x$ | The regressor in the bias regression function. |
| $\beta$ | The regression coefficients in the bias regression function. |

The same assumptions made in [22] are followed.

- Each transcript is independently processed, then sequenced. Given the parent trail-scripts, reads are independently generated.
- the probability of observing a read is read type *s*_*j*_ of length *l*_*i,j*_ does not depend on the transcript and is equal to the empirical insert length distribution *q*(*l*_*i,j*_).
- Given a read insert length *l*_*i*,*j*_ and transcript of origin, the start site distribution *t*(*ψ*_*i*_,_*j*_) is uniform and its support is all the possible positions this read can be generated from.
- The random variable *n*_*i*,*j*_, the number of reads from read type *s*_*j*_ that are generated from transcript *i*, follows a Poisson distribution with parameter *θ*_*i*_*a*_*i,j*_.

The insert length model defines the probability of a read of type *s*_*j*_ observed after processing transcript *i* as:

#### Definition 4

*Pr*(*a read r of type s_j_ observed after sequencing transcript i*) = *Pr*(*r ∈ s_j_*|*r ∈ i*) *Pr*(*r ∈ i*)

where

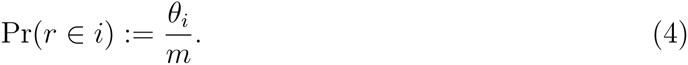

The probability of selecting a isoform in a given locus is approximated as the fraction of isoform *i* in the nucleotide space [13]. As opposed to the model of [22], our parameterization does not involve the isoform length.

#### Definition 5

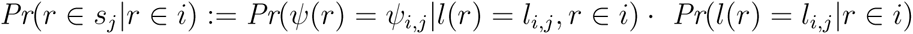

The probability of observing a read of read type *s*_*j*_ given that the isoform which generates this read is *i*, is equal to the product of two conditional probabilities. i.e. the probability of observing a read's insert length given its parent transcript and the probability of observing the 5'end of the read given both its isoform of origin and insert size. These two probabilities distributions are known as insert size distribution and start site distribution.

Thus we can write the joint probability as

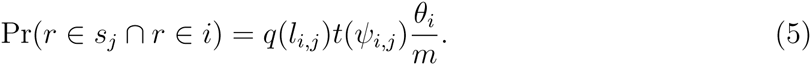

The random variable *n*_*i*,*j*_, the number of reads from read type *s*_*j*_ that are generated from transcript *i*, follows a Poisson distribution with mean expressed as:

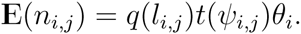

The model assumes that each *n*_*i*,*j*_ is independent. And since the sum of independent Poisson distributed random variables is a Poisson random variable, we have

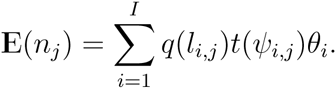

and summing over reads on exon bin *k* leads to,

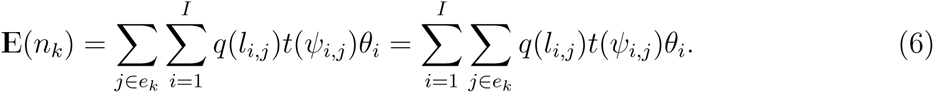

Therefore the parameter vector Θ is linked to the observation vector [*n*_1_,*n*_2_,…,*n*_*K*_] by the9 Poisson mean parameters. In a more compact way and using sampling rate notation, equation 6 can be written as

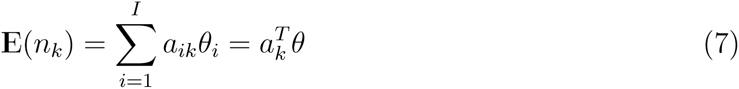

Here, *a*_*ik*_ = Σ_*j*∈*ek*_*q*(*l*_*i*,*j*_)*t*(*ψ*_*i*,*j*_). Calculating *a*_*ik*_ requires summing over all the possible read types over an exon bin *k*. Since *t*(·) is a uniform distribution, we can instead sum over all the possible insert lengths.

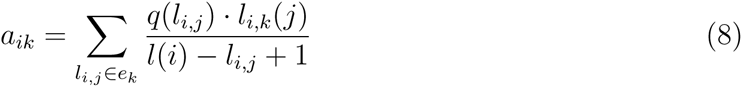

*l*_*i*,*k*_(*j*) is the number of possible positions where a read type *s*_*j*_ can be generated on the *i*th isoform and *k*th exon bin. However, calculating *l*_*i*,*k*_(*j*) is not trivial since it depends on the number of exons and their length in the exon bin, read length and gap length. Its computational complexity grows exponentially as the number of exons of a exon bin grows 7.1. Luckily, the number of exons belonging to a exon bin is usually not very large. We observed at most 5 exons for a exon bin in our simulation.

In a follow up study of *Jiang and Salzman* [10], the authors proposed adding a auxiliary bias parameter to each of the Poisson counting units, i.e. in their case, *n*_*j*_ and our case *n*_*k*_. This model is over parametrized and they have to use regularization to make it justifiable. Strawberry, instead, uses a linear regression to calculate *b*_*k*_ in a such way

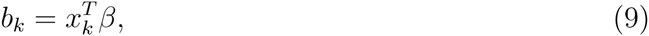

where *β* is the regression coefficients. Numerous studies have shown that bias is cause by locus sequences (e.g., hexamer bias) around the reads [8], position of the reads[21], GC content bias[2], etc. Therefore we believe there is a pattern in the bias and want to use a simple linear regression to capture the pattern. The simple linear bias model can be extended to incorporate further bias pattern yet to be discovered. In the current implementation, three regressors, bin length and bin GC content and average insert size are included. Under the assumption that the bias is multiplicative as did in [10], we have a new Poisson mean as the product of 7 and 9,

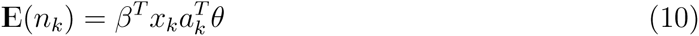

Equation 10 indicates an identity link function for generalized Poisson regression. Poisson regression usually uses log as the link function to avoid the logarithm of a negative number when optimizing over log likelihood function. However since all the numbers are positive in our case, we can avoid the negative value issue using the EM algorithm [4]. Finally, the log likelihood function for this identity link Poisson regression is:

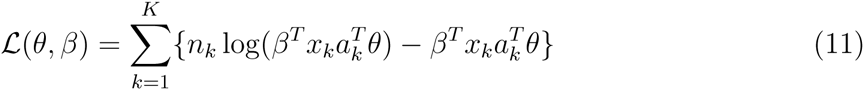

This is a bi-concave function in which for fixed *β*, function 11 is concave in *θ* and for fixed *θ*, it is concave in *β*. *θ* and *β* are estimated using the technique in [10], called Alternative Concave Search(ACS), which alternatively fixes one and estimates the other until convergence. In our implementation, we use the EM algorithm for maximizing *θ* and *β* iteratively while holding the other constant at each time.

#### Algorithm 2

With β fixed, θ can be solved with the following EM algorithm.

- E-step:

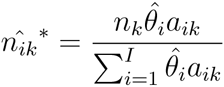
- M-step:

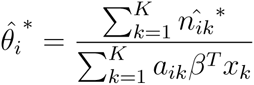

With θ fixed, β can be solved with using the EM algorithm in a similar fashion.

- E-step:

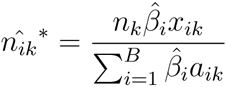
- M-step:

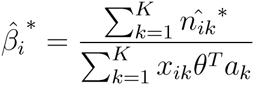

Here, i indexes the bias regressors.

The EM algorithm also provides an intuition for conducting bias correction. When updating θ, the bias term only affects the M step of that EM algorithm. This can be seen as a reweighting strategy. Without the bias term, 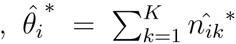 and 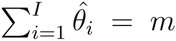. The algorithm assigns exactly *m* reads to different isoforms. Under the influence of bias, each bin gets weighted up if it is underrepresented or weighted down if overrepresented in therms of the number of reads. This eliminates the implicit constraint that the sum of *θ̂*_*i*_ is *m*.

### 5.5 Empirical insert length distribution and converting half-mapped reads to concordantly mapped reads

Before starting the estimation procedure, Strawberry calculates the empirical insert length distribution. This is done by looking at the concordantly mapped reads on loci where only a single isoform is present after assembly step. To fully utilize the insert length and orientation information from the read alignment, Strawberry converts half-mapped reads (only one end of a read pair is actually mapped) to fully mapped paired-end read by probabilistically generating the other end. To simulate a paired-end read from an half-mapped read, a insert length is simulated from a truncated normal distribution *N*(*μ*,*σ*). *μ* and *σ* are empirical distribution mean and standard variance respectively and the normal distribution is truncated at the minimum and maximum insert length. According to the orientation and location of the existing end the location of absent end can be calculated. If the new end extends beyond its transcript boundary it will be truncated exactly at the boundary. If the distance from the existing end to the transcript boundary is less than the minimum insert length observed this read will be discarded since it is very likely to be a false alignment. In our simulated paired-end RNA-seq dataset[15], 20% are only half-mapped using TopHat2 [11].

## 6 Acknowledgements

The material presented here is based upon work supported by the National Science Foundation under Grant IOS-1062546. Any opinions, findings, and conclusions or recommendations expressed in this material are those of the author(s) and do not necessarily reflect the views of the National Science Foundation.

## 7 Appendix

### 7.1 Number of possible positions for a read to start on an exon bin

For a bin that contains at most 2 exons, it is straightforward to compute the number of possible starting positions for a read type*j*, isoform *i* and bin *k*, i.e., *l*_*i*,*k*_(*j*) as:

- Bin size 1: *l*_*i*,*k*_(*j*) = *l*(*k*) – *l*_*i*,*j*_ + 1
- Bin size 2: *l*_*i*,*k*_(*j*) = min(*l*_left_, *l*_*i*,*j*_ – 1) + min(*l*_right_, *l*_*i*,*j*_ – 1) – *l*_*i*,*j*_ + 1

*l*_left_ is the length of the leftmost exon and *l*_right_ is the length of the rightmost exon. When the size is larger than 2, we can partition the number of positions for a read to hit at least both ends of the bins, denoted by *T*, to 2^*s*−1^ classes. Here, to hit means only the sequenced ends lie within certain exons and *s* is the number of exons in the bin. In the bin size 3, for example, (Fig. 4(b)), there are *T*_13_ number of positions for a read hitting *e*1 and *e*3 but not *e*2 and *T*_123_ number of positions for a read hitting all three exons. *T* = *T*_13_ + *T*_123_. We have two utility functions to calculate *T*_13_ and *T* in constant time. Thus *T*_123_ is obtained by subtracting *T*_13_ from *T*. For a exon bin of size of 4 or more, *l*_*i*,*k*_(*j*) can be solved in a recursive way by using these two utility functions.

~~~
l _ all <-function (l _ left, l _ right, l _ int, r, gap){
   #return number of possible postions such that read hits
   #at least farthermost exons in the bin.
   #r, the read length
   #gap, the gap length
   #l_left, the leftmost exon length of the bin
   #l_right, the rightmost exon length of the bin
   #l_int, the total length of intern exons of the bin

   L = 2*r +gap – l _ int – 1;
   return (min(l_left, L) + min(l_right, L) – L)
}
l_gap <-function (l _ left, l _ right, l _ int, r, gap){
   #return number of possible positions such that read hits #only farthermost exons of the bin, not any internal exons. low = max(r, l _ left + l _int - gap - 1);
   up = min(l_left, l _ left + l _ right + l _ int –gap – r); return (max(0, up - low))
}
~~~

### 7.2 Details about using CuffCompare

We extracted only multi-isoform genes from Arabidopsis TAIR10 model and then ran Cuffcompare to compare the assembly results from all three programs to this reference.

~~~
cuffcompare –r ref.g t f assembled_transcripts.gtf
~~~

Two outputs, the file with suffix *.stat* and the file with suffix *.tmap* from Cuffcompare were further processed for comparing assembly accuracy and quantification accuracy respectively.

